# A Versatile GPMV-Imaging Platform for Quantitative Analysis of Receptor Binding and Membrane Fusion

**DOI:** 10.1101/2025.05.29.656808

**Authors:** Inbar Yosibash, Suman Khan, Alisa Vaknin, Raviv Dharan, Alexandra Lichtenstein, Susan Daniel, Ori Avinoam, Raya Sorkin

## Abstract

Membrane fusion is central to biological processes such as viral entry, fertilization and cell-to-cell fusion. Gaining a mechanistic understanding of fusion requires the ability to visualize and quantify the dynamic interaction between two membranes and their associated protein machineries at high temporal and spatial resolution. However, studying these processes in live cells remains challenging due to the complexity of the cellular environment. Here we demonstrate a versatile cell-free platform based on giant plasma membrane vesicles (GPMVs) that enables controlled, quantitative analysis of receptor binding and membrane fusion kinetics in a native membrane context. As proof of concept, we reconstitute the SARS-CoV-1 Spike-ACE2 interaction, capturing specific receptor engagement and accumulation at the membrane interface using confocal microscopy and micropipette aspiration. Fusion was induced by proteolytic activation and quantified using both high-resolution microscopy and high-throughput Imaging Flow Cytometry. The platform also reveals the influence of membrane composition on fusion efficiency, demonstrated by the impact of cholesterol depletion. This approach provides a broadly applicable system for dissecting membrane fusion and protein-protein interactions across membranes, with compatibility for biophysical, imaging and structural analysis. It offers new opportunities for mechanistic studies and inhibitor screening in a biologically relevant yet experimentally accessible context.

## Introduction

Membrane fusion is the process by which two distinct lipid bilayers combine to form a single, continuous membrane. This is a fundamental cellular process essential for various functions such as viral entry, intracellular trafficking, and developmental events such as fertilization and myogenesis ^1-3^. Enveloped viruses, including Coronaviruses, exploit membrane fusion to enter host cells by fusing their lipid envelope with host cell membranes ^4, 5^. This process is driven by specialized membrane proteins called fusogens, which interact with cell-surface receptors and undergo regulated conformational changes that catalyze the energetically unfavorable fusion reaction ^6^.

In Coronaviruses such as SARS-CoV-1 (CoV-1), membrane fusion is mediated by the Spike glycoprotein, a class I fusogen with distinct structural domains: S1, which binds to the ACE2 receptor, and S2, which contains the conserved fusion machinery ^7, 8^. Binding of Spike to ACE2 initiates a cascade of conformational changes in the S2 domain, typically following proteolytic cleavage, bringing the two membranes into close apposition and driving their merger ^4, 9, 10^. Depending on the availability of host proteases, fusion may occur directly at the plasma membrane or following endocytic uptake. Despite the importance of this mechanism, the intermediate stages of receptor binding, protein clustering, membrane rearrangements, hemifusion, and pore expansion remain difficult to study in a physiologically relevant yet experimentally tractable way.

Dissecting the sequence and regulation of these steps requires a platform that provides spatiotemporal resolution, native membrane composition, and compatibility with quantitative imaging and biophysical measurements. To reveal mechanistic details of fusion processes, it is also advantageous to use single vesicles with natively embedded membrane proteins and to capture large numbers of interactions. Current experimental approaches face limitations. Liposome-based assays lack the lipid and protein complexity of native membranes and require biochemical reconstitution of functional proteins. Supported bilayers can introduce artifacts due to membrane immobilization. Live cell approaches, while physiologically relevant, are limited by both the crowded nature of cell cultures and the complexity of the intracellular environment, making it difficult to capture individual fusion events, as their location and timing are unpredictable and challenging to visualize in three dimensions. As a result, it has remained difficult to deconstruct the biophysical and molecular contributions of receptor binding and membrane organization leading to membrane fusion in a controlled and accessible system.

Here, we present a versatile, high-throughput and simple Giant Plasma Membrane Vesicles (GPMVs)-based platform that bridges this gap. GPMVs are cell-derived vesicles that preserve close to native lipid and protein composition of the parent cell plasma membrane while eliminating confounding intracellular processes and the actin cortex ^11^. We combined GPMVs with high-resolution live imaging, micropipette aspiration, and high-throughput Imaging Flow Cytometry to analyze membrane fusion and receptor binding kinetics in a controlled, cell-free context. As proof of principle, we used the SARS-CoV-1 Spike-human ACE2 interaction. We visualized specific receptor clustering at the membrane interface and quantified fusion efficiency after proteolytic activation of the Spike. We further validated the sensitivity of the assay by showing that cholesterol depletion significantly reduces fusion, consistent with the known role of cholesterol in viral entry ^12-14^.

Compared to existing approaches, our GPMV-based platform uniquely enables the quantitative, biophysical dissection of protein-protein interactions and membrane fusion across native membranes. It is readily adaptable to other fusogen-receptor pairs and can support a range of imaging, biophysical and structural biology techniques, including cryoEM. By offering both mechanistic insight and the experimental flexibility of an *in-vitro* system, this approach opens new avenues for studying membrane fusion in systems ranging from viral entry to developmental cell-cell fusion and for screening inhibitors in a biologically meaningful setting.

## Results

### Forming GPMVs with labeled content or proteins

To establish a simplified, biologically relevant platform for studying protein-protein interactions and membrane fusion, we generated GPMVs with defined proteins of interest. As a proof of concept, we chose the well characterized interaction between the SARS-CoV1 Spike fusogen and its host cell receptor, ACE2.

We generated two distinct populations of GPMVs derived from mammalian cells. One population expressed ACE2 fused to GFP, allowing visualization of receptor distribution in the membrane. The second population expressed Spike, with cytoplasmic mCherry as a content marker (Fig. 1 *A*; see Materials for details, Supplementary movie 1). Importantly, labeling the Spike GMPVs via cytoplasmatic expression of mCherry avoided direct modification of the fusogen itself, minimizing the risk of perturbing its activity.

**Figure 1.**
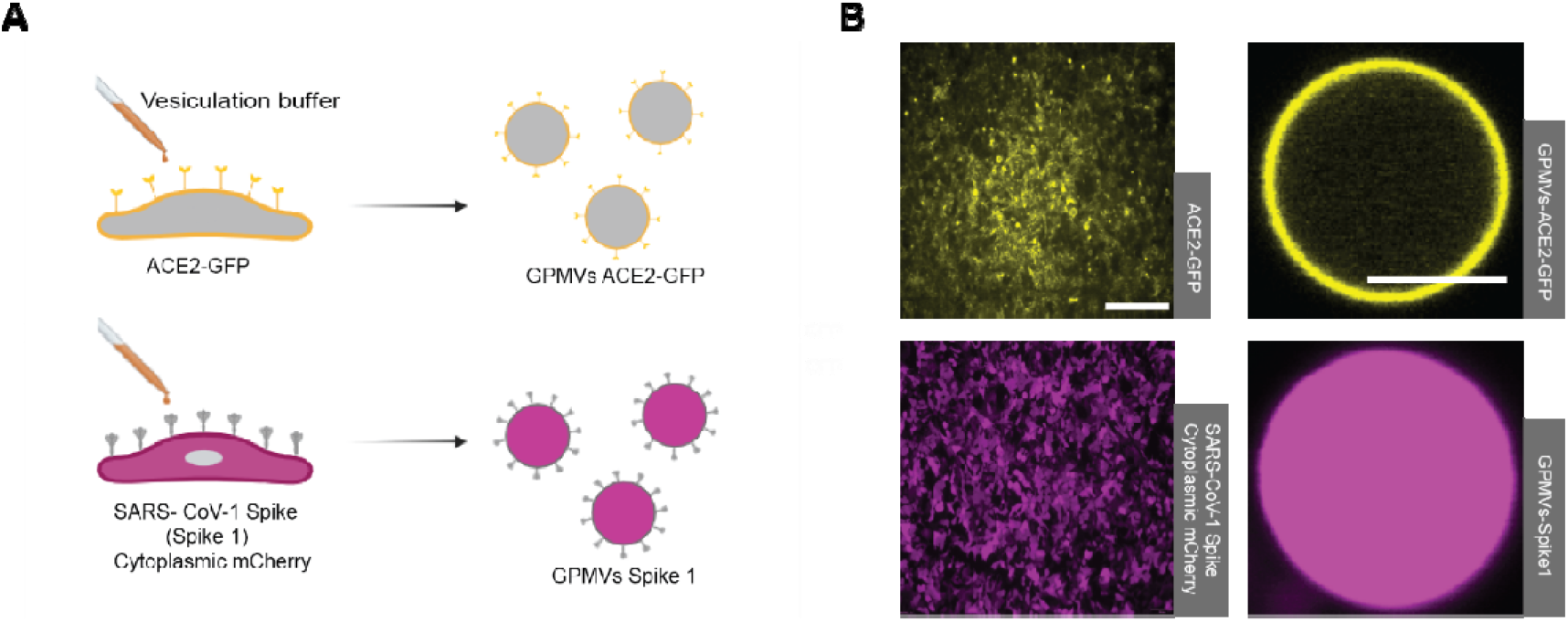
The formation of GPMVs: (A) Schematic illustration of GPMV formation. (B) Microscopy images of Flp-In TREx 293 cells expressing ACE2-GFP/ HEK293 cells expressing Spike and cytoplasmic mCherry (left Images), scale bar 500µm. The isolated GPMVs contained Spike in their membrane and cytoplasmic mCherry/ ACE2-GFP in their membrane (right Images), scale bar 5 µm.

Fluorescence microscopy of the cell cultures confirmed the robust expression and correct localization of the fluorescent proteins in the parent cells and in the GPMVs. ACE2-GFP was incorporated into the GPMV membranes, while mCherry was retained in the lumen of Spike expressing vesicles, yielding two clearly distinguishable vesicle populations (Fig. 1 *B*). This configuration enabled simultaneous monitoring of membrane-membrane interactions via ACE2 monitoring and content mixing via mCherry under controlled conditions. The ability to manipulate and track each membrane population independently provided a unique opportunity for kinetic analysis of receptor engagement, as well as quantitative assessment of fusion events.

### Receptor binding and docking dynamics in the GPMV Platform

To assess the capability of our GPMV-based system to capture early events in membrane fusion, we first investigated the interaction between ACE2 and SARS-CoV-1 Spike, focusing on the initial receptor engagement and docking dynamics. GPMVs bearing ACE2-GFP and those containing Spike-mCherry were mixed in the presence of 2⍰mM Ca^2+^ (Fig. 2 *A*). Time-lapse confocal microscopy revealed that upon mixing, ACE2-GFP–positive GPMVs rapidly adhered to Spike-expressing GPMVs, with visible enrichment of ACE2-GFP fluorescence at the membrane contact site (Fig. 2 *B*). This spatial distribution suggests receptor clustering or accumulation at the fusion interface prior to fusion.

**Figure 2.**
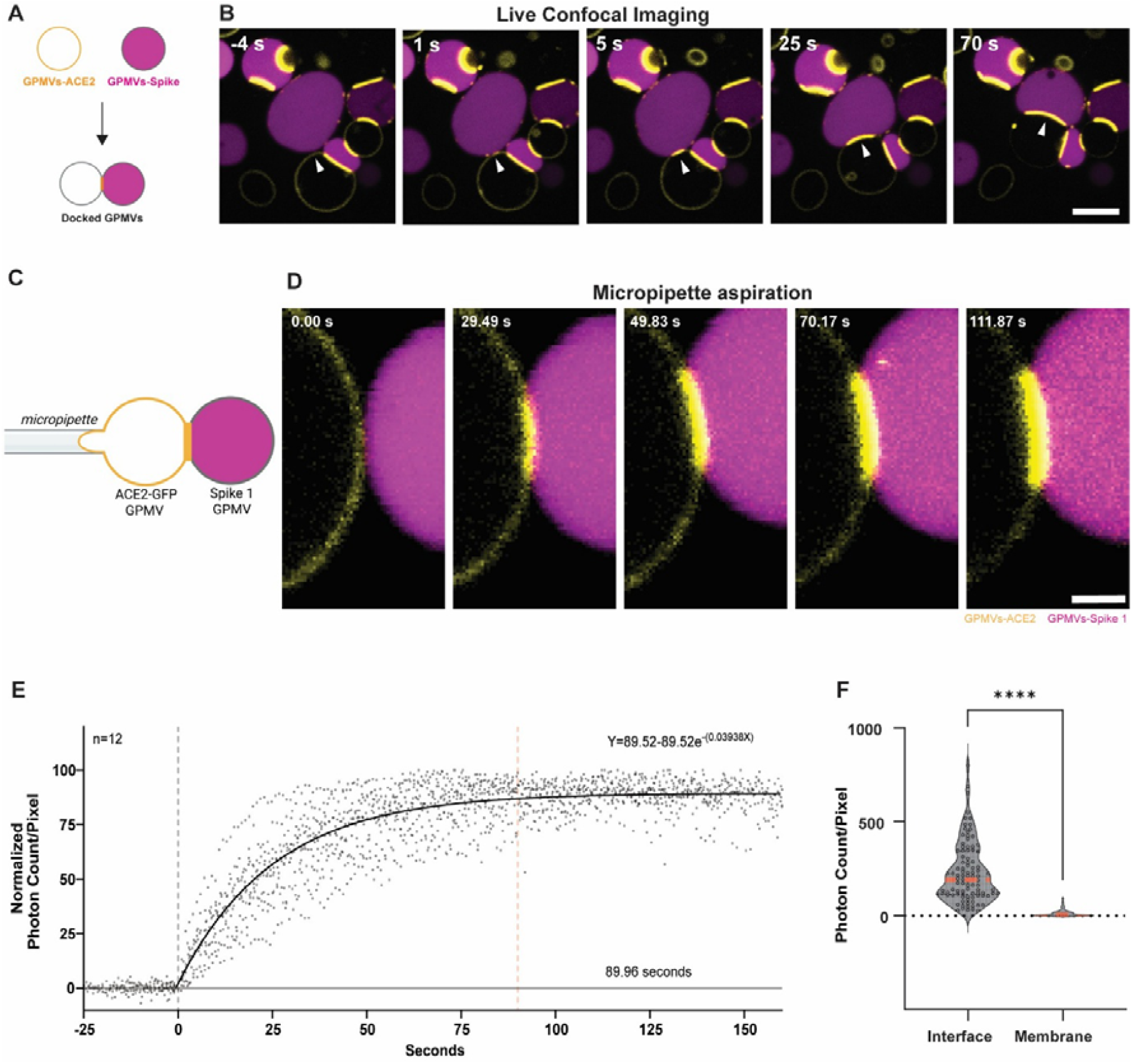
Visualization of receptor binding between Spike and ACE2 GPMVs: (A) Schematic illustration of the experimental procedure of GPMV docking. (B) Time-lapse live confocal microscopy imaging of GPMV docking. Both types of GPMVs mixed with 2mM Ca^2+^ and their docking dynamics were observed. The white arrowheads indicate a representative docking event. Time stamps (in seconds) denote the progression of docking, scale bar 10µm. (C) Schematic illustration of the experimental setup of (D). (D) Time-lapse live confocal microscopy imaging of GPMV docking using the C-trap confocal microscope and micropipette aspiration. A GPMV expressing ACE2-GFP (yellow) was trapped using a micropipette and brought close to a Spike-expressing GPMV (magenta) to observe docking dynamics. The time stamps (in seconds) indicate the progression from initial contact to docking, scale bar 2µm. (E) Kinetics of reaching docking steady state. Exponential fitting of GPMV docking kinetics measured using live confocal microscopy and micropipette aspiration. The plot shows normalized photon counts per pixel over time, representing ACE2-GFP accumulation at the docking interface. The black curve represents an exponential fit (n = 12 vesicles). The docking process reached a steady state at ∼90 seconds (indicated by the red dashed line), where the grey dashed line determines time zero. (F) Violin plot comparing the photon counts per pixel of the interface and the ACE2-GFP membrane of the docked GPMVs (n = 53 vesicles). The black dots represent experiments conducted, and the red line represents the median (P < 0.0001, Mann-Whitney).

To quantify the kinetics of this interaction, we used a micropipette-based assay to bring individual GPMVs into contact under controlled conditions (Fig. 2 *C*). This enabled precise initiation of receptor-ligand engagement and real-time observation of ACE2 accumulation at the interface. Fluorescence intensity increased over time and reached equilibrium at ∼90 seconds (n = 12, Fig. 2 D and *E*, Supplementary movie 2), indicating rapid equilibration of receptor binding and recruitment.

To evaluate whether this behavior was influenced by vesicle immobilization, we performed parallel experiments with freely diffusing vesicles, tracking spontaneous docking events in solution using confocal microscopy. These exhibited a comparable equilibration time of 85.4 seconds (n = 23, Supplementary figures 1 and 2, Supplementary movie 3), suggesting that vesicle mobility does not significantly affect the intrinsic kinetics of ACE2-Spike binding in this system.

To assess the degree of receptor accumulation at the interface, we compared the photon counts per pixel of ACE2-GFP in the docking region to its baseline membrane levels in freely moving, undocked, vesicles. We observed a statistically significant enrichment of ACE2-GFP fluorescence at the docking interface (P < 0.0001, n = 53, Fig. 2 *F*), indicating receptor redistribution at the contact zone.

Together, these results validate our platform as a sensitive, quantitative system for monitoring receptor-ligand interactions across native membranes. The ability to compare free and immobilized vesicle interactions, coupled with precise temporal resolution, enables detailed dissection of binding kinetics and spatial receptor organization, which are key early steps in fusogen-mediated membrane fusion.

### Triggering and quantifying membrane fusion using GPMVs

To test whether full membrane fusion can be reconstituted and observed in our GPMV system, we asked whether proteolytic activation of the Spike would trigger fusion between docked vesicles. Previous studies have shown that trypsin cleavage of the Spike protein promotes the conformational changes required for membrane merger ^15-17^. We therefore mixed ACE2-GFP and Spike mCherry GPMVs to allow docking, then added trypsin to induce fusion (Fig. 3 *A*).

**Figure 3.**
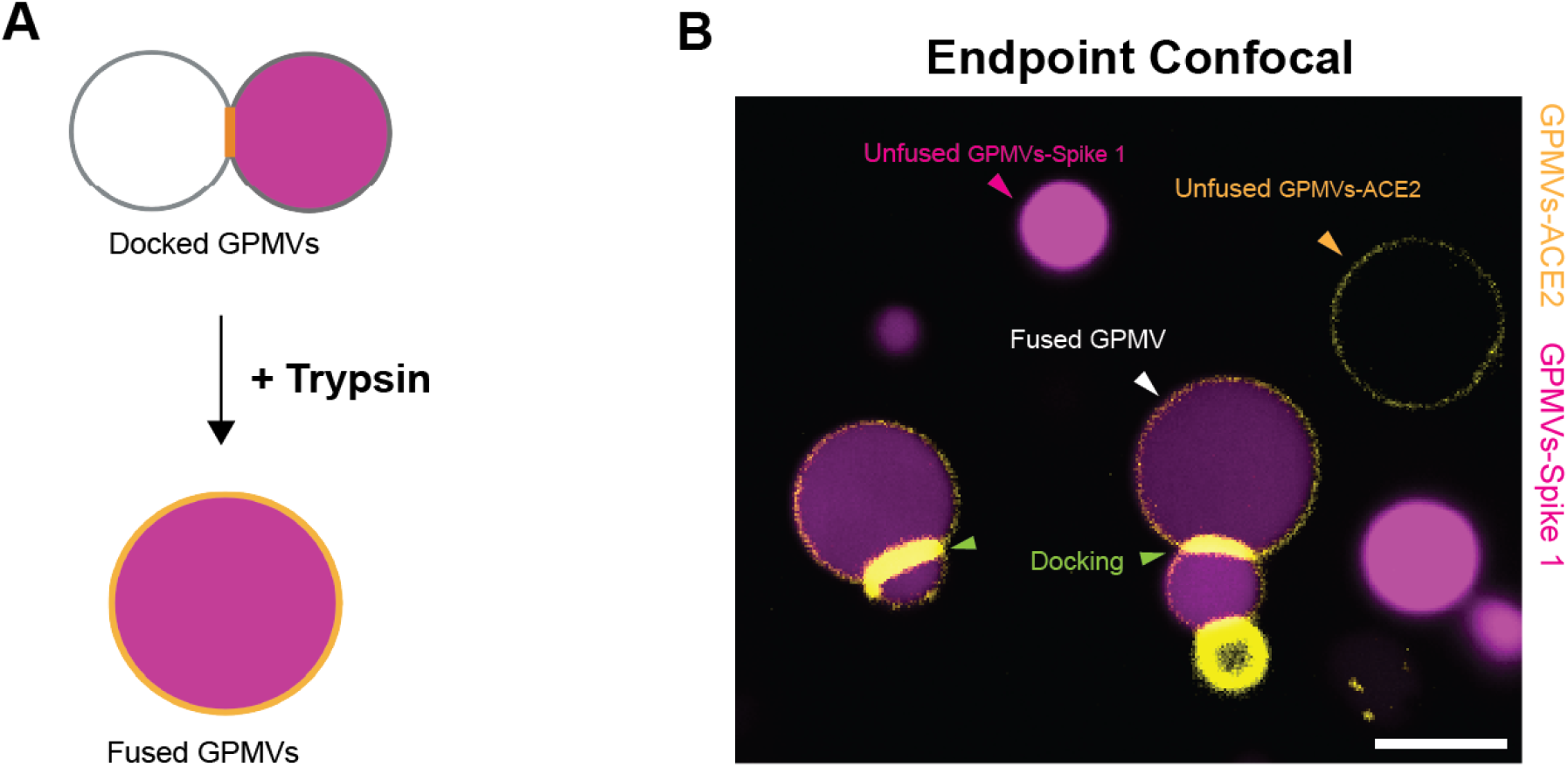
Visualization of trypsin-triggered Spike-ACE2 GPMV full fusion in real time. (A) Schematic illustration of the experimental procedure of GPMVs fusion. (B) Endpoint states. Representative field including unfused Spike with mCherry GPMV (magenta arrowhead), unfused ACE2-GFP GPMV (orange arrowhead), fused GPMV (white arrowhead), and GPMV docking (green arrowhead), scale bar 10µm.

Live confocal microscopy revealed the occurrence of full fusion events, seen as the mixing of mCherry from Spike GPMVs into ACE2-GFP vesicles (Fig. 3 *B*). Time-lapse microscopy demonstrated that fusion occurred rapidly after protease treatment, with most observable events taking place within the first three minutes (Supplementary movie 4 and 5). This dynamic visualization of fusion in real time highlights the ability of the platform to capture temporal features of the fusion process in a controlled environment.

To quantify fusion efficiency, we acquired four fields of view per experiment, each containing an average of ∼159 vesicles, and measured the fusion index two hours after trypsin addition. The fusion index was defined as the percentage of vesicles containing both GFP and mCherry fluorescence relative to the less abundant vesicle population. We observed a consistent fusion index of ∼30±5 (N = 3), indicating robust and reproducible fusion under these conditions (Fig. 3 *B*).

Notably, vesicles that had undergone fusion were still capable of docking and fusing with additional vesicles, as shown by the presence of multiple labeled vesicles engaged in further membrane contacts. This observation suggests that fused vesicles retain fusogenic potential, allowing for analysis of sequential or cumulative fusion events in future studies.

### High Throughput quantification of fusion using Imaging Flow Cytometry

While confocal microscopy provides high-resolution insights into vesicle fusion events, it has limitations. The presence of out-of-focus vesicles complicates image analysis, and the inherently low-throughput nature of the technique restricts the number of vesicles that can be quantitatively assessed. To overcome these limitations, we turned to Imaging Flow Cytometry (ImageStream), which combines high-throughput sampling with single vesicle image acquisition, enabling the detection and classification of thousands of GPMVs in suspension without the spatial constraints of traditional microscopy

We performed ImageStream analysis two hours after trypsin treatment and observed that unfused single vesicles and fused GPMVs are readily distinguishable (Fig. 4 *A,* supplementary Fig. 3). We obtained a fusion index of ∼33±9 (N = 7), in close agreement with our confocal microscopy-based measurements (Fig. 4B). This consistency between modalities validates the robustness of the assay and supports the use of ImageStream for large-scale fusion screens.

**Figure 4.**
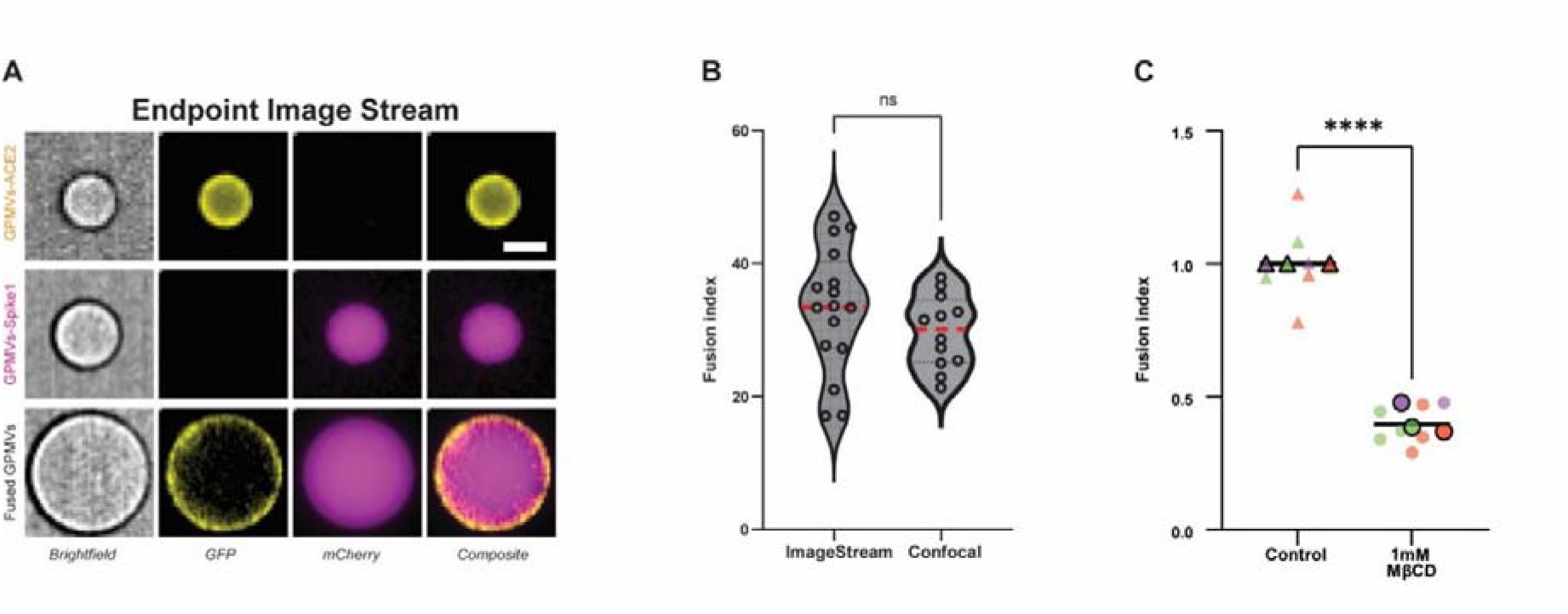
ImageStream can assess fusion efficiency and identify differences under cholesterol depletion. (A) ImageStream Imaging Flow Cytometry GPMVs fusion endpoint (i.e., 2 hours after triggering). Representative images show brightfield, mCherry (yellow), GFP (magenta), and composite channels for individual GPMVs and fused vesicles. Top row displays GPMV expressing ACE2-GFP (yellow), middle row displays GPMV expressing Spike (labeled with mCherry) and bottom row displays a fused GPMV showing the colocalization of GFP and mCherry signals in the composite image, scale bar 5µm. (B) Comparison of fusion index (%) between ImageStream and confocal microscopy. The fusion index was calculated as the number of GPMVs showing colocalization of mCherry and GFP divided by the less abundant GPMV population and multiplied by 100. Data are shown for ImageStream fusion index (∼33±9, N = 7) and confocal microscopy (∼30±5, N = 3). The black dots represent experiments conducted, and the red line represents the median. (C) Cells expressing ACE2-GFP and cells expressing Spike-mCherry were treated with 1mM methyl-β - cyclodextrin for 30 minutes prior to GPMV generation. Fusion index was quantified using ImageStream, and normalized to untreated controls. The black line represents the overall mean, and each outlined shape indicates the mean of a single experiment. Cholesterol depletion resulted in a significant reduction in fusion (P < 0.001, Mann-Whitney test). Colors correspond to independent experiments.

To test the sensitivity of the platform to perturbations, we examined the effect of plasma membrane cholesterol depletion, a known modulator of viral membrane fusion. Treatment of both GPMV populations with 1mM methyl-β-cyclodextrin, as previously reported for SARS-CoV-1 fusion ^12^, led to a significant ∼2.5-fold reduction in fusion efficiency compared to controls (P < 0.001, Fig. 4 *C*). These results underscore the requirement of cholesterol for efficient fusion in this system and further establish the GPMV-based assay as a platform that faithfully recapitulates a physiologically relevant membrane parameter.

Altogether, the combination of confocal microscopy and Imaging Flow Cytometry provides a powerful and scalable framework for studying receptor-mediated membrane fusion, enabling both mechanistic dissection and high-throughput screening in a biologically relevant yet experimentally tractable setting.

## Discussion

Membrane fusion is essential in many physiological and pathological processes, including viral infection and fertilization ^1-3^. Due to its importance, various experimental methods have been developed over the years to study fusion. Here, we introduce a new method to investigate membrane fusion and protein-protein interactions in a controlled, cell-free, yet biologically relevant environment using GPMVs.

To demonstrate that GPMVs can be effectively used to test fusion, we used the well-established membrane fusion reaction mediated by the viral Spike protein and the human ACE2 receptor as a model. To characterize the ACE2-Spike interactions, we tracked in real time the accumulation of ACE2-GFP at the interface between GPMVs using confocal fluorescence microscopy within our optical tweezers instrument. The accumulation kinetics reached a steady state within approximately 90 seconds (Fig. 2 *E*), suggesting a rapid and specific interaction between Spike and ACE2. To assess binding specificity, we used GPMVs expressing the membrane protein TSPAN4 conjugated to GFP, and observed no docking events under identical conditions (Supplementary Fig. 4). The kinetics were similar with micropipette-immobilized vesicles and the freely diffusing vesicles (Supplementary Fig. 2), confirming that surface interactions did not impact the binding between ACE2 and Spike at these conditions. This result demonstrates that the GPMV system is a faithful model for studying early events in viral entry.

To quantitatively study fusion, we imaged GPMVs following their mixing and fusion-triggering using confocal fluorescence microscopy. Confocal microscopy provides high-resolution, real-time insights into the fusion process (Fig. 3 *B*). Additionally, it enables various real-time manipulations, such as photoactivation and photobleaching experiments. However, it presented certain limitations: the presence of out-of-plane vesicles complicated accurate quantification as it was often difficult to determine whether vesicles observed in the focal plane were part of a fusion event or simply overlapping, and the low-throughput nature of confocal imaging limited the number of vesicles that could be analyzed in a reasonable time frame, reducing statistical power. In each experiment, we captured four fields of view, with an average of 159 vesicles per field. To measure a larger population of free-standing, non-supported vesicles, we expanded the use of Imaging Flow Cytometry (ImageStream), a technique commonly used for extracellular vesicles (EVs) and synthetic vesicles. ImageStream enables the simultaneous collection of high-resolution images and flow cytometric data from thousands of individual vesicles allowing sensitive detection of single vesicles, morphological characterization, and the ability to distinguish true vesicle fusion events from vesicle overlap or background noise ^18, 19^. This dual capability is especially valuable when working with heterogeneous vesicle populations, where accurate classification is critical ^20, 21^. ImageStream has also been shown to detect differences in vesicle size, shape, and fluorescence intensity ^22^, and to detect different EVs in complex biological fluids such as blood plasma ^23^

We mixed ACE2 harboring GPMVs with a population of GPMVs with activated spike protein, and screened the vesicles using flow cytometry (Fig. 4 *A,* Supplementary Fig 3). ImageStream flow cytometry is highly suitable for our assay due to its high-throughput capability, enabling us to analyze approximately 5,000 vesicles within 10 minutes. The fusion index obtained using confocal microscopy was ∼30±5 and ∼33±9 in ImageStream (Fig. 4 *B*). The similarity between ImageStream and confocal microscopy results further emphasized the applicability of ImageStream for quantitative fusion studies. In our hands, ImageStream effectively distinguished fused GPMVs from single vesicles two hours post-trypsin treatment, complementing the lower-throughput, high-resolution data obtained by confocal microscopy.

To test whether differences in fusion efficiency can be detected with ImageStream, we depleted plasma membrane cholesterol using 1mM methyl-β-cyclodextrin (MβCD), a water-soluble oligosaccharide composed of β(1–4)-linked glucopyranose units arranged in a closed ring structure. Its external surface is hydrophilic, while its inner cavity is hydrophobic. This structure enables MβCD to enhance the solubility of hydrophobic compounds, such as cholesterol, by encapsulating them within its inner cavity ^24^. Cholesterol depletion disrupts membrane microdomains enriched in cholesterol ^12, 25^ which are critical for viral entry processes. Consistent with previous studies ^12-14^, We observed an approximately 2.5-fold reduction in fusion efficiency following cholesterol depletion (Fig. 4 *C*).

Several experimental approaches have been commonly used to study membrane fusion. Liposome ensemble assays rely on bulk fluorescence measurements, typically using dequenching to monitor lipid and content mixing between vesicles ^26^. These assays provide a quantitative ensemble measure of fusion, however, they average over large populations of vesicles, cannot resolve individual fusion events or intermediates, and are prone to artifacts from leakage and non-linear quenching ^27^. Single-vesicle fusion assays allow real-time observation, such as kinetics of single vesicle fusion events ^28, 29^, and are often based on large or small unilamellar vesicles (LUVs and SUVs). This assay includes vesicle fusion to supported planar membranes or to other immobilized vesicles. Fluorescent labeling enables the detection of fusion intermediates using fluorescence microscopy. However, interactions with the support can hinder membrane dynamics, and the small size of the vesicles makes it difficult to control membrane tension ^27^. On larger scales, giant unilamellar vesicles (GUVs) fusion assays enable direct visualization of fusion events ^30^ and morphological changes using optical and electron microscopy ^31, 32^. Although fusion between GUVs permits observation of lipid and content mixing as well as mechanical manipulation of membrane tension, reconstituting fusion proteins in GUVs is technically challenging ^27^. Another common approach is to use whole cells ^33^. While this preserves native membrane complexity, studying fusion mechanisms in live cells is challenging due to their complex composition and topography ^11^. However, our system has limitations: GPMVs lack cytoskeletal structure and active machinery, which can be necessary for certain fusion events in live cells. Importantly, both interacting membranes are of the same size and exhibit negligible curvature, unlike viral membranes, which have high curvature that can influence fusion ^34^. Additionally, vesicle size heterogeneity and variations in protein expression levels may introduce some variability in efficiency and kinetics. Future improvements, such as using size-sorted vesicles and methods to achieve a more uniform protein expression, could further improve the reproducibility and precision of this assay.

Our GPMV-based platform offers a powerful tool for studying membrane fusion in a biologically relevant yet simplified environment. Beyond viral entry, this approach could be applied to investigate fundamental mechanisms of membrane fusion and screen potential inhibitors. Future studies could expand this system to other fusion proteins and environmental conditions, further broadening its applicability in membrane biology and therapeutic development.

## Materials and Methods

### Cell Culture

HEK293T (ATCC CRL-3216) or Flp-In T-REx 293 stable ACE2-GFP cells were cultured in DMEM medium (Sartorius 01-055-1A) containing 10% Fetal bovine serum (Gibco A5256701), 2mM glutamate (Sartorius 03-020-1A), 1mM sodium pyruvate (Sartorius 03-042-1B) and 1% penicillin– streptomycin (Gibco 15140122) in 100mm cell culture plates (Corning) at 37°C and 5% CO_2_. Dissociation of the cells was achieved with 1mL of trypsin-EDTA (Bio-Lab 004715233100). Cells were collected after centrifugation at 100g for 3 minutes, diluted 1:20, plated on a 100mm plate and cultured until the confluency reached 50−60% and no more than 24 hours before transfection.

### Plasmid Preparation and Stable Cell Lines Generation

The plasmid encoding SARS-CoV-1 Spike, including its map is provided in the Supplementary (Supplementary Fig. 5). The plasmid encoding mCherry was obtained from Clontech (catalog no. 632523). Plasmids scale-up was performed using Nucleobond midi-plasmid purification kit (Macherey-Nagel no. 740410). The sequence was verified by Sanger sequencing (ZABAM Instrumentation and Service, Tel Aviv University).

The hACE2-eGFP gene (Addgene plasmid # 154962) was subcloned into pcDNA™5/FRT/TO (Thermo Fisher Scientific) using NEBuilder® HiFi DNA Assembly (NEB) following the manufacturer’s protocol. The primers (IDT) used for the PCR amplification were:

Insert - 5’ AACTTAAGCTTGGTACCGAGGCTAGCGTTTAAACGGGC 3’ and 5’ ACTGGACTAGTGGATCCGAGTTACTTGTACAGCTCGTCC 3’,

Vector - 5’ CTCGGATCCACTAGTCCAG 3’ and 5’ CTCGGTACCAAGCTTAAG 3’.

The eGFP-pcDNA™5/FRT/TO plasmid was a gift from Dr. Ariel Ben-Bassat (Tel Aviv University). The cloned gene sequences were verified by Sanger sequencing (ZABAM Instrumentation and Service, Tel Aviv University). Stable cell lines were generated by Lipofectamine™ 2000-mediated transfection of ∼1 million Flp-In™ T-REx™ 293 cells (Thermo Fischer Scientific) plated in a 6-well plate (Corning) with 2.7µg of pOG44 (Thermo Fisher Scientific) and 0.3µg of gene of interest containing pcDNA™5/FRT/TO constructs. Cells were maintained at 37°C, 5% CO_2_ in Dulbecco’s Modified Eagle Medium (Sartorius) supplemented with 10% FBS, 2mM L-Glutamine (Thermo Fischer Scientific), and 1mM Sodium Pyruvate (Sartorius). For stable selection, the media was supplemented with 50µg/mL hygromycin and 15µg/mL blasticidin (Invivogen) three days post-transfection, with hygromycin increased to 100µg/mL after seven days. Additionally, 100µg/mL Penicillin and 100µg/mL Streptomycin were added to maintain sterility. Media was replaced every 2–3 days until sufficient cell islands formed, after which cells were expanded for cryopreservation.

### HEK293T Cell Transfection and Flp-In T-REx 293 Cell Induction

HEK293T or Flp-In T-REx 293 stable ACE2-GFP cells were plated at 20% confluency on a 100mm plate coated with 10μg/mL Poly-L-lysine (Sigma-Aldrich) to keep the cells attached and to minimize cell debris in solution. At 50% confluency, HEK293 cells were transiently cotransfected with 12μg mCherry plasmid and 12μg Spike plasmid using 60μl Lipofectamine 2000 (Invitrogen, Thermo Fisher scientific) according to the manufacture’s protocols. Flp-In T-REx 293 stable ACE2-GFP cells were induced by adding 1μg/mL tetracycline (Sigma-Aldrich) to the medium. The cells were grown at 37°C and 5% CO_2_ for 24 to 36 hours to obtain an optimal protein expression.

### Depletion of Cholesterol in Cells Prior to GPMV Production

To deplete cholesterol, 100 mm plates containing cells expressing either Spike and mCherry or ACE2-GFP were washed twice with 10 mL PBS. Then, 1 mM Methyl-β-cyclodextrin (Sigma-Aldrich, C4555) was added to the plates, followed by incubation for 30 minutes at 37°C and 5% CO_2_. In parallel, control cells (expressing either Spike-mCherry or ACE2-GFP) were incubated in DMEM without MβCD for the same duration. Both cell populations—Spike-mCherry–expressing cells and ACE2-GFP–expressing cells—underwent cholesterol depletion prior to GPMV production for the fusion experiments. For control experiments, GPMVs were prepared from cells treated with DMEM only.

### GPMVs Formation

Following protein expression, 100mm plates with cells were washed with 10mL GPMV buffer (20mM HEPES and 150mM NaCl at pH 7.4). GPMV vesiculation was induced using a published methods ^35^. Briefly, the cells were washed and incubated with 4mL active vesiculation buffer (GPMV buffer supplemented with 2mM CaCl_2_, 1.9mM dithiothreitol (DTT) and 27.6mM formaldehyde) at room temperature for at least 4 hours in the dark, during which GPMVs are spontaneously formed and released into the supernatant. The GPMVs were collected using a cut pipette tip. A centrifugation step of 100g for 3 minutes was employed to remove any cell debris, then, the supernatant was collected. The GPMVs were used on the same day.

### Sample Preparation for ImageStream and Confocal Fluorescence experiments

For all experimental setups, GPMVs expressing ACE2-GFP and GPMVs expressing Spike-mCherry were mixed in the presence of 2 mM CaCl_2_ and incubated at 37°C and 5% CO_2_ for at least 2 hours, allowing membrane docking to occur.

Docking Assays: After incubation, samples were directly used for docking analysis.

ImageStream Assays: Following the 2-hour docking incubation, samples were further incubated for 2 hours with 0.83 g/L trypsin (Sigma-Aldrich, T1426) at 37°C and 5% CO_2_ prior to imaging.

Docking Kinetics (C-trap Confocal): The mixed GPMVs were immediately imaged following the addition of CaCl_2_, and docking kinetics was monitored in real time.

### Control experiments with TSPAN4-GFP GPMVs

HEK293T cells were seeded at 20% confluency on 100 mm plates pre-coated with 10⍰μg/mL poly-L-lysine (Sigma-Aldrich, P6282) to ensure cell adhesion and reduce the presence of cell debris in solution. Upon reaching ∼50% confluency, cells were transiently transfected with 10⍰μg of a TSPAN4-GFP expression plasmid (Dharan et al., 2022) using 20⍰μL of Lipofectamine 2000 (Invitrogen, Thermo Fisher Scientific), and incubated at 37°C in a 5% CO_2_ atmosphere for 24–36 hours to allow for protein expression. GPMVs were subsequently prepared as described above. For docking experiments, TSPAN4-GFP-expressing GPMVs were mixed with GPMVs expressing Spike fused to cytoplasmic mCherry in the presence of 2 mM CaCl_2_, and incubated at 37°C with 5% CO_2_ for at least 2 hours.

### Confocal and Micropipette Aspiration Setup

The docking kinetics experiment was performed using a C-trap confocal fluorescence optical tweezers setup (LUMICKS) made of an inverted microscope based on a 60X water-immersion objective (NA 1.2). A micropipette aspiration setup including a micromanipulator (Sensapex) holding a micropipette with a diameter of 5μm (Biological industries) was connected to a Fluigent EZ-25 pump and integrated into the optical tweezer instrument. Before and after each experiment, the zero-suction pressure was found by aspirating a polystyrene bead (3.43μm, Spherotech) into the pipette and reducing the suction pressure until the bead stopped moving. Using micropipette aspiration, one ACE2-GFP GPMV was trapped and brought into proximity with Spike mCherry GPMV while tracking the time until reaching the steady state of the docking. The micropipette pressure used was less than 0.1mbar and therefore the GPMV tension was not affected. Experiments were performed at room temperature.

### Spinning Disk Confocal Microscopy

Live receptor binding and fusion experiments were performed using a Yokogawa automatic Spinning Disk confocal scanning unit (CSU-W1-T2) mounted on an inverted Olympus IX83 microscope (37°C, with 5% CO_2_). 60× 1.4 NA oil immersion objectives were used for data acquisition. Images were captured by back-illuminated Prime 95B sCMOS cameras (Photometrics) controlled by VisiView software (Visitron Systems GmbH). Confocal fluorescence excitation was done with solid-state laser diodes (488nm for eGFP and 561nm for mCherry). The following fluorescence emission filters were used: 525/50nm for eGFP and 609/54nm for mCherry.

### Intensity Analysis at Membrane Interfaces of Immobilized GPMVs

A custom graphical user interface (GUI) was developed to assist manual analysis of fluorescence intensity distributions at membrane interfaces from time-series microscopy images. Raw TIFF sequences were first pre-processed using percentile-based contrast normalization for visualization while preserving the original bit depth for quantitative analysis.

To distinguish between interface and non-interface membrane regions with precise spatial control, a dual region of interest (ROI) approach was implemented where two rotatable rectangular regions were manually positioned on each frame: (1) Interface ROI, placed directly at the membrane contact site to quantify local enrichment; and (2) Reference ROI, positioned at a non-interface membrane region to establish baseline intensity values from the same image. Each ROI could be freely rotated and repositioned with sub-pixel accuracy to accommodate membrane geometries ensuring accurate tracking of membrane interfaces throughout the time series.

Mean fluorescence intensities were calculated for both interface and reference regions at each time point by averaging all pixel values within each mask. For time-series analysis, ROI configurations could be propagated across frames with position correction capabilities to maintain consistent region definitions throughout the sequence. Results were automatically exported as CSV files containing time points and corresponding mean intensity values for both regions.

### Intensity Analysis at Membrane Interfaces of Moving GPMVs

To quantify intensity distributions at membrane interfaces from time-series fluorescence microscopy data of free-floating GPMVs, we integrated deep learning-based segmentation with quantitative intensity profiling.

Two-channel TIFF stacks were first preprocessed with percentile-based contrast normalization for segmentation and tracking while preserving the original bit depth for quantitative analysis. For vesicle tracking, reference points on both vesicles of interest within the same frame were manually defined. These points served as input to a SAM2 (Segment Anything Model 2) neural network which propagated segmentation masks across all frames, enabling robust tracking and segmentation of freely moving and deformable GPMVs throughout the time series.

To analyze the membrane interface, we applied computational geometry to detect intersection regions between vesicles, generating perpendicular profile lines that extended from the interface into both vesicles. This automated approach eliminates the both bais and labor-intensive nature of manual ROI placement in our previous method, while providing more precise localization of interface boundaries even as GPMVs pair move and rotate.

Membrane-specific “donut” masks were created through adaptive boundary detection based on intensity gradients at membrane peripheries. The algorithm automatically identified membrane boundaries by analyzing radial intensity profiles, where values exceeding 75% of peak intensity defined the inner membrane boundary, while the outer boundary was established at 80% of peak intensity along the outward gradient. This adaptive approach ensures consistent isolation of membrane regions despite variations in membrane thickness or fluorescence intensity.

Intensity measurements were calculated for both interface and non-interface membrane regions by extracting pixel values along the profiling lines within the donut masks. The analysis was exported as CSV files containing time points and corresponding intensity values for both regions.

### IMultispectral Imaging Flow Cytometry (ImageStreamX)

GPMVs were prepared and mixed as described above and imaged using multispectral imaging flow cytometry (ImageStreamX markII flow cytometer; Cytek Biosciences). Single and focused population of GPMVs was gated using brightfield channel: Gradient RMS (feature measures image contrast or focus quality) and morphological features (area and aspect ratio of GPMVs). Between 4 and 8 thousand events of single focused GPMVs were gathered for each sample and visualized. eGFP fluorescence was measured using 488nm wavelength laser,200mW for excitation and channel 2 band emission spectrum of 480-560nm. mCherry fluorescence was measured using laser 561nm wavelength, 200mW for excitation and channel 4 band emission spectrum of 595-642nm. Experiments were performed at room temperature.

### ImageStream Fusion Quantification

To quantify the number of fused GPMVs obtained from ImageStream, all image channels (brightfield, GFP, and mCherry) were first normalized to 300 pixels along the largest dimension while preserving the aspect ratio with Hermite interpolation (Supplementary Fig. 6). Following this pre-processing, all image channels were segmented using the Cellpose neural network to generate instance masks of the GPMVs in all three channels.

To identify spherical GPMVs from cellular membrane aggregates and debris, a weighted sphericity assessment was performed on each GPMV instance mask obtained from the BF channel as reference. The following circularity metrics and weights were used: (1) Perimeter-Area ^36^ Circularity (35% weight)— implementing the isoperimetric quotient (4π·area/perimeter^2^) to identify shapes approaching perfect circularity; (2) Distance Transform Circularity (35% weight) — quantifying radial uniformity by calculating the variation coefficient of centroid-to-boundary distances and transforming it via exponential function (e^(-2·variation)) to penalize irregularities; (3) Convexity-Based Circularity (15% weight) — measuring the area-to-convex-hull-area ratio to identify concavities that deviate from ideal vesicle morphology; and (4) Eccentricity-Based Circularity (15% weight) — calculated as (1-eccentricity) to account for elongation. Values greater than 0.71 were used to select round GPMVs.

To determine whether each reference circular GPMV was eGFP-positive, mCherry-positive, or both, cross-channel colocalization analysis was performed pairwise (i.e., BF-GFP and BF-mCherry masks), requiring mask area overlap (≥70%) with respect to the reference channel. Based on this cross-channel colocalization, GPMVs were classified as either eGFP-positive, mCherry-positive, or both, while GPMVs lacking eGFP or mCherry signal were discarded. The fusion index was calculated as 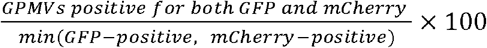, using the minimum channel positivity count as the denominator to compensate for potential variations in fluorophore labeling efficiency between markers.

### Fusion Index Normalization

Fusion index values were normalized to the mean of the control group. For each experiment, the mean of the control replicates was calculated, and both control and treated (1mM methyl-β-cyclodextrin) values were divided by this average. This normalization set the control group to a mean of 1, allowing the treated group to be expressed as fold change relative to control. Normalized values were used for analysis and visualization.

### Docking Normalization and Quantification

To quantify the ACE2-eGFP accumulation at the docking interface, fluorescence values were normalized where 0% represented the intensity before the start of interaction and 100% corresponded to the intensity when steady state was reached, following the formula: normalized value = [(original value - minimum value) / (maximum value - minimum value)] × 100. For each interacting pair, the start of accumulation was manually identified and set as the zero time point for analysis. The normalized data were then fitted to characterize the accumulation kinetics and determine the time to equilibrium using the exponential plateau model: Y = YM - (YM - Y0) × exp(-K × X), where Y0 is the starting fluorescence intensity at the manually aligned starting point, YM is the maximum fluorescence intensity at steady state, K is the rate constant (inverse of time units), and X is time. Both normalization and curve fitting were performed using GraphPad Prism built-in functions.

### Methyl-β-cyclodextrin Versus Control Normalization

Raw fusion index values were normalized to the average of the control group within each experiment. Specifically, the mean fusion index of the control samples was calculated, and all corresponding control and 1mM treatment values were divided by this mean.

## Code Availability

Code used for the data analysis is available at:

https://gitlab.com/sorkin-lab/gpmv-fusion-project

## Acknowledgements

I.Y. acknowledges Financial support from the ADAMA Center. We thank Dr. Ziv Porat, Head of the Flow Cytometry Unit at the Weizmann Institute of Science, for valuable discussions regarding the ImageStream experiments. R.S. acknowledges support by the Israel Science Foundation (Grant No. 1289/20), and the NSF-BSF (Grant No. 2021793). Co-Funded by the European Union (ERC ReMembrane 101077502). Views and opinions expressed are however those of the authors only and do not necessarily reflect those of the European Union or the European Research Council Executive Agency. Neither the European Union nor the granting authority can be held responsible for them. O.A. acknowledges funding from the Abisch-Frenkel Foundation, the Minerva Foundation with funding from the Federal German Ministry for Education and Research and the Minna James Heineman Foundation, the Henry Chanoch Krenter Institute for Biomedical Imaging and Genomics, the Schwartz Reisman Collaborative Science Program, and the Yeda-Sela Center for Basic Research. O.A. is an incumbent of the Miriam Berman presidential development chair.

