## Supplementary material for "A Versatile GPMV-Imaging Platform for Quantitative Analysis of Receptor Binding and Membrane Fusion": SI

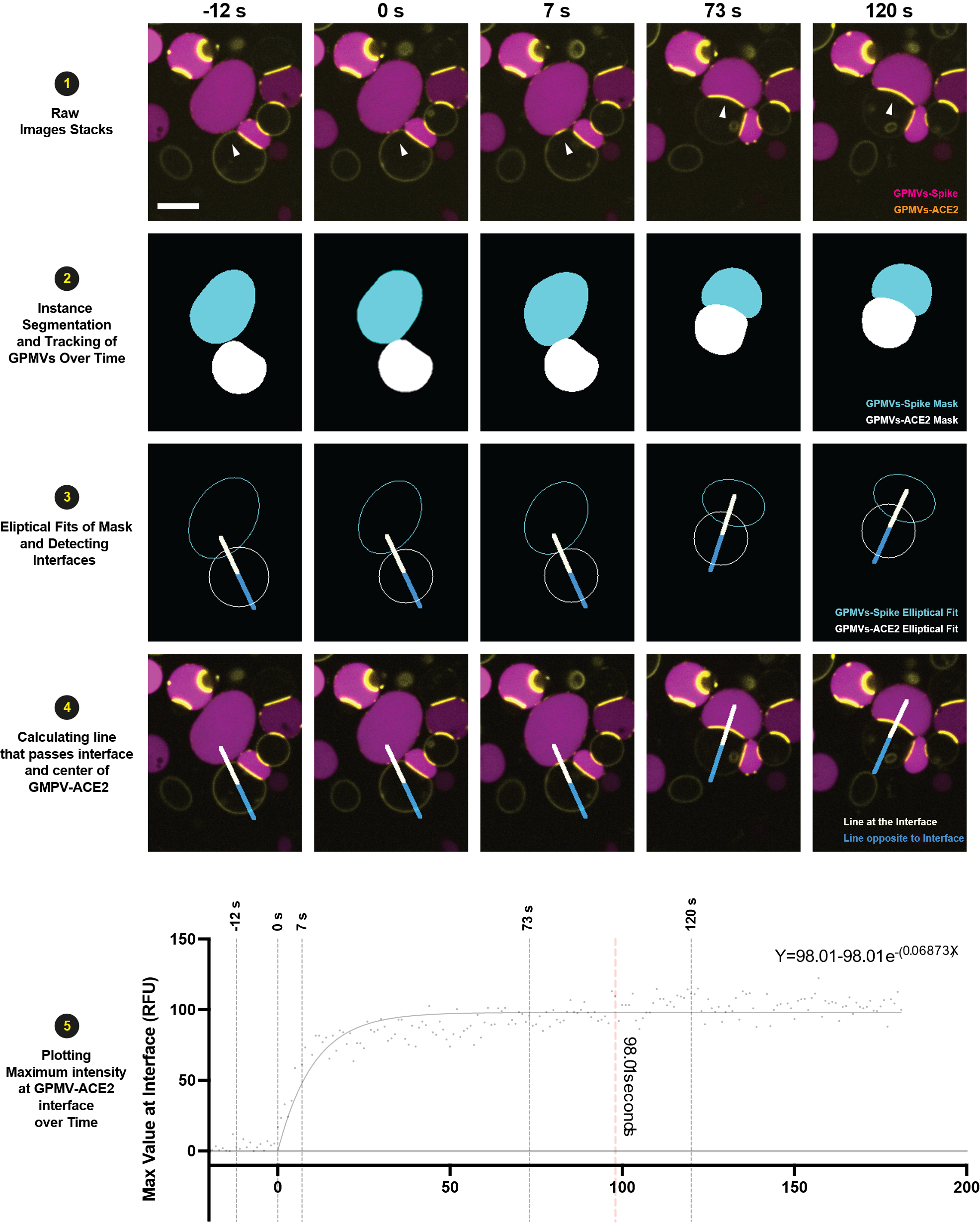

**Supplementary Figure 1: An artificial intelligence-based analysis pipeline to quantify protein-protein interactions.** Our image analysis pipeline quantifies protein interactions at GPMV interfaces by live confocal microscopy (1). Using artificial intelligence-based tracking, we track vesicle movement and identify membrane boundaries (2) and subsequently the contact interface with a line (3,4). We generate time-resolved fluorescence intensity profiles across these interfaces (4,5), quantifying protein accumulation and docking kinetics during GPMV interactions (5). The graph shows exponential fitting of ACE2-GFP accumulation at the docking interface. The black curve indicates the exponential fit, with steady state reached at ~98 seconds (red dashed line). Grey dashed lines mark corresponding time points shown in the frames above.

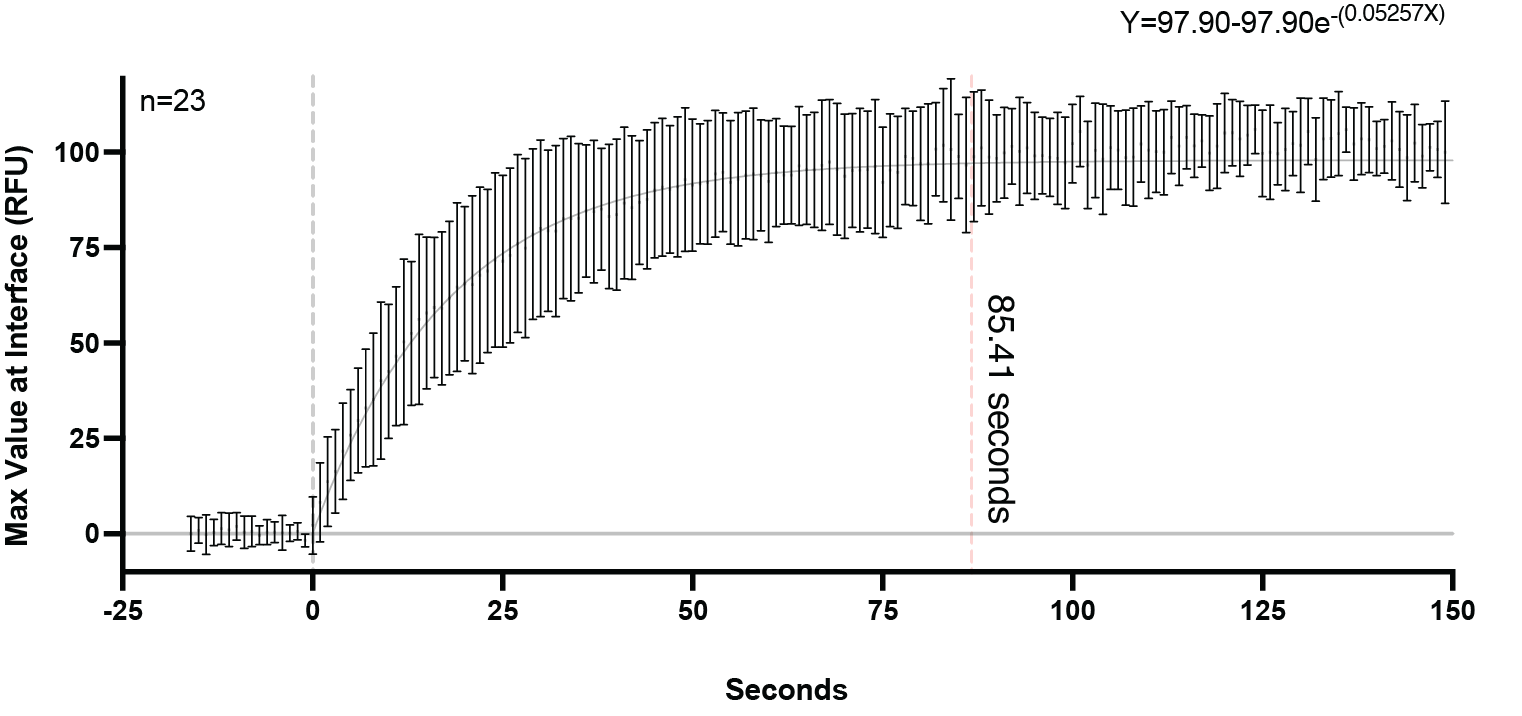

**Supplementary Figure 2:** **Kinetics of docking between ACE2-GFP and Spike 1 using live confocal imaging**. The plot shows exponential fitting of ACE2-GFP accumulation at the docking interface across 23 events where both GPMVs were mobile. The black curve indicates the exponential fit, with steady state reached at ~85 seconds (red dashed line), comparable to results observed in immobilized vesicle pairs (Fig 2C & F).

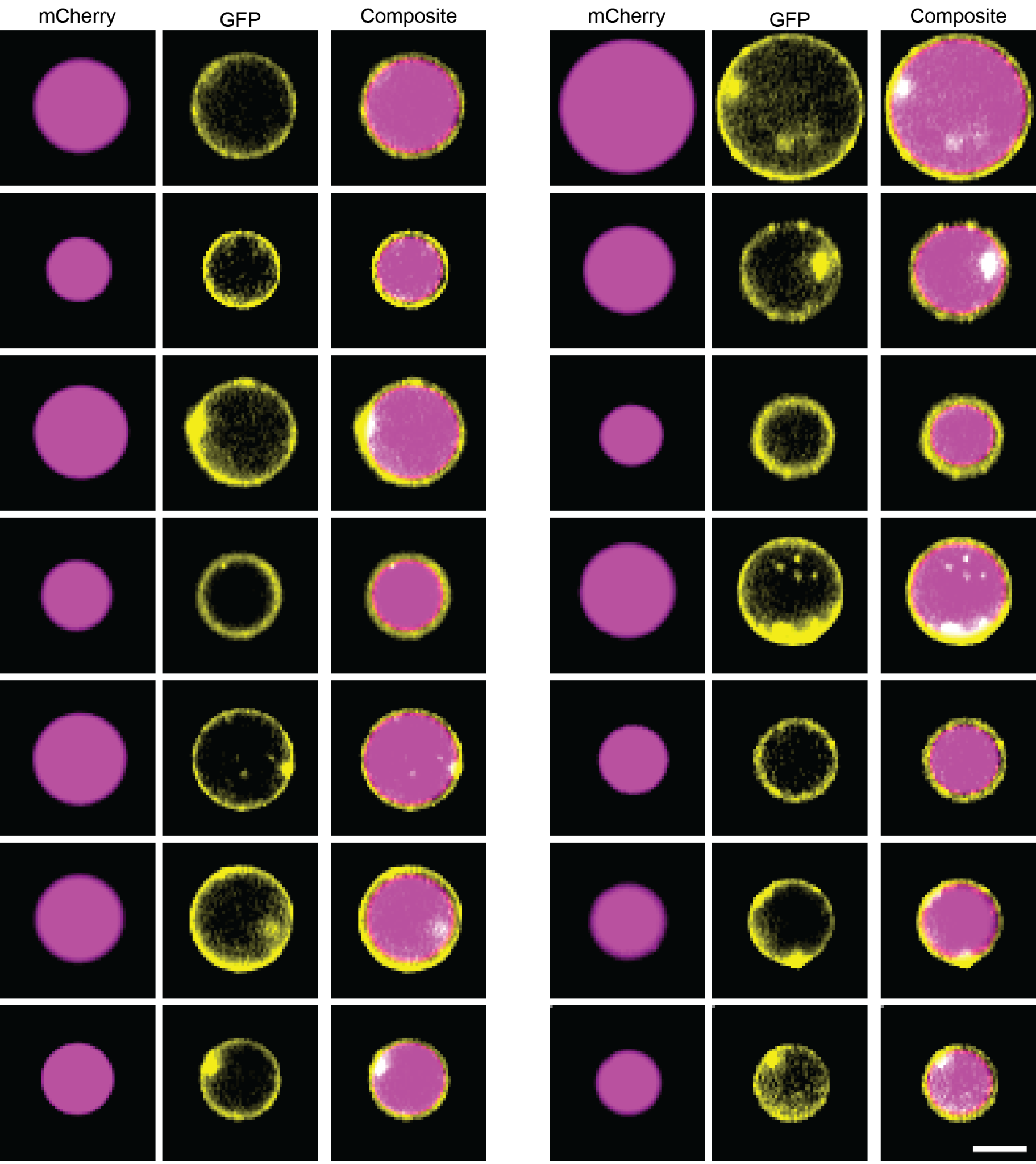

**Supplementary Figure 3:** ***ImageStream enables visualization of fused GPMVs.*** *ImageStream images of GPMVs fusion endpoint. mCherry (yellow), GFP (magenta), and composite channels for fused vesicles. Scale bar 5um.*

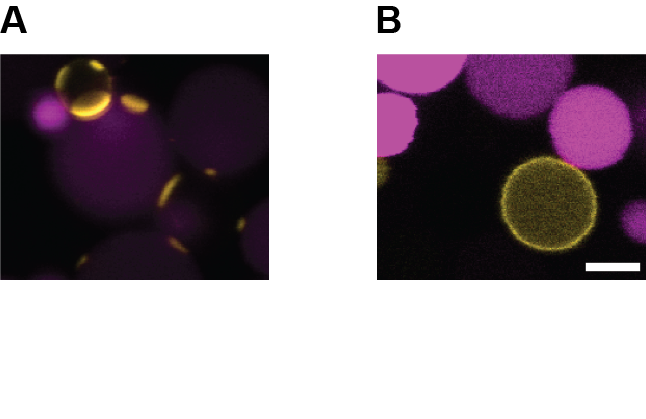

**Supplementary Fig. 4: Specific interactions of Spike- ACE2.** (A) GPMVs expressing Spike (magenta) and ACE2- GFP (yellow) were mixed in the presence of 2mM Ca^2+^ and incubated at 37°C and 5% CO_2_ for 2 hours. Docking was observed in all fields of view. (B) GPMVs expressing Spike (magenta) and TSPN4-GFP (yellow) were mixed in the presence of2mM Ca^2^ and incubated at 37°C and 5% CO_2_ for 2 hours. No docking was observed in any of the frames*.* Scale bar 5µm.

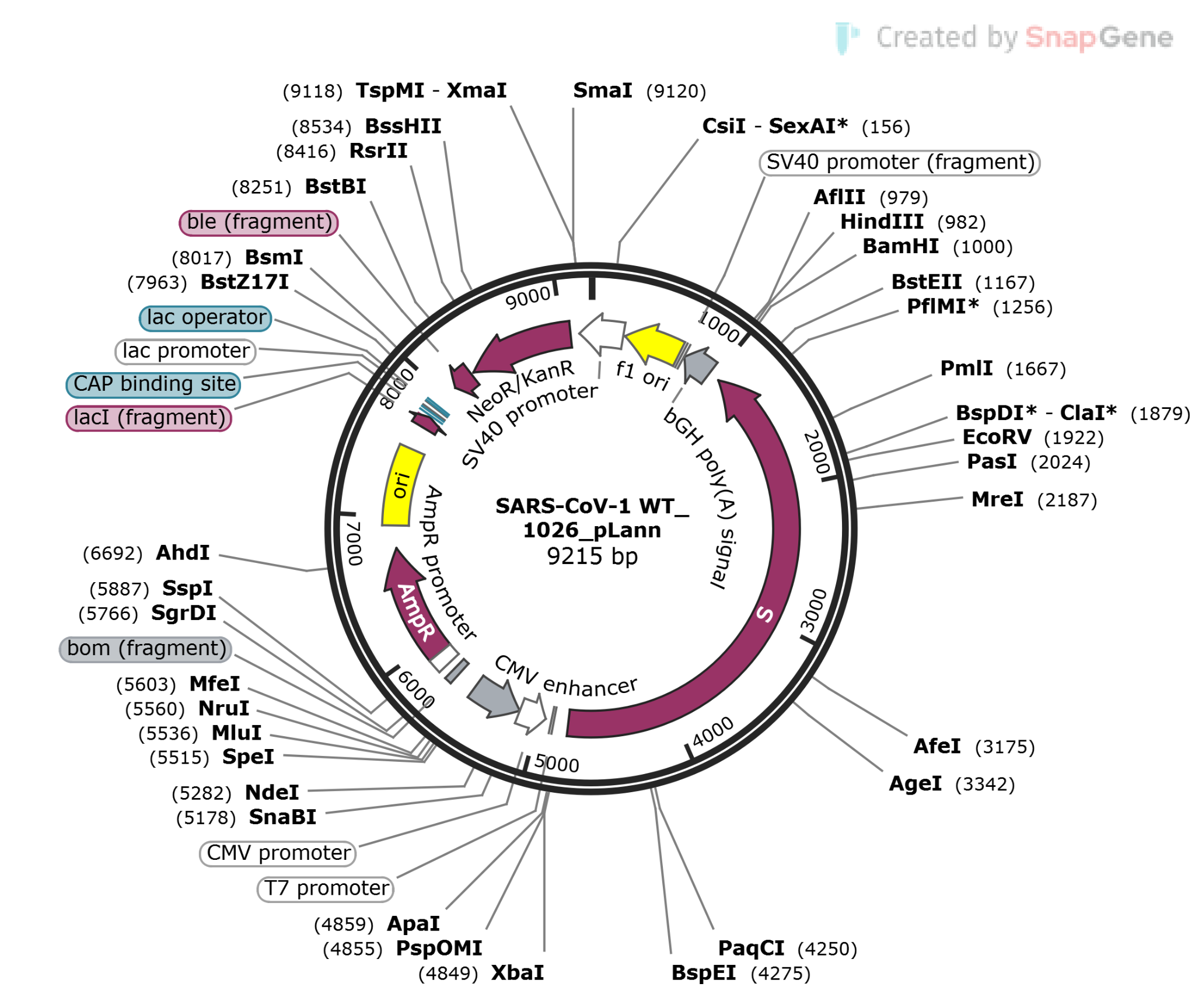

**Supplementary Figure 5:** Plasmid map of SARS-CoV-1 WT_1026_pLann (9215 bp), used for expression of SARS-CoV-1 Spike (S) protein.

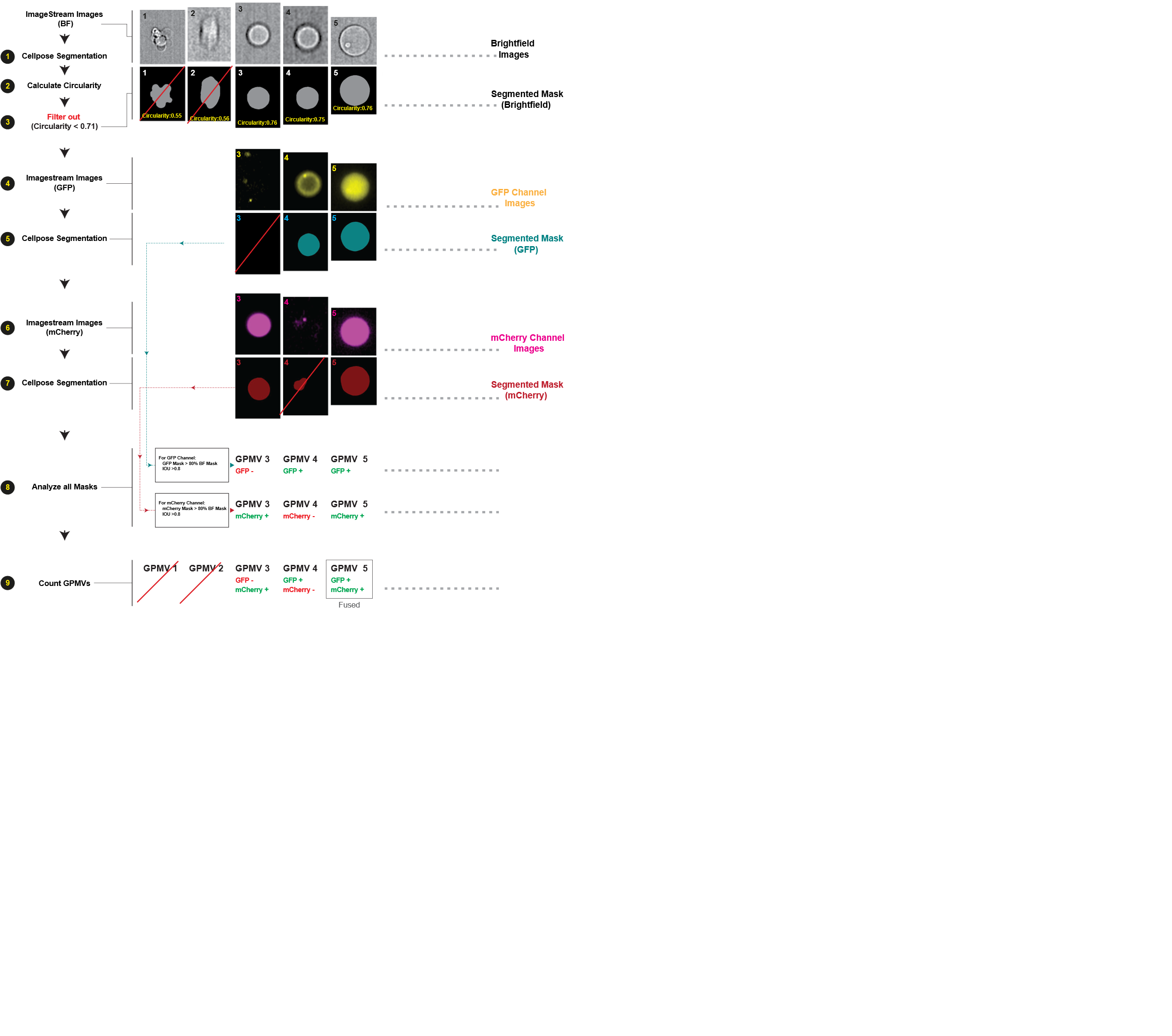

**Supplementary Figure 6:** **Quantification of GPMV fusion using neural network segmentation**. We take brightfield (BF) channel images (1), apply Cellpose segmentation (2), and filter by circularity (threshold >0.71) to isolate true GPMVs from membrane fragments (3). We then process GFP channel images (4) with Cellpose segmentation (5), and repeat for mCherry channel (6-7). We identify fluorescently-labeled vesicles by requiring >80% overlap between each fluorescence mask and the brightfield mask (8). We then classify vesicles as GFP-positive, mCherry-positive, or dual-positive (fused) (9) and calculate the fusion index as the percentage of dual-positive GPMVs relative to the minimum count of single-labeled populations.

**Python modules used in the study**

**Image Processing and Computer Vision:** 'PIL', 'tifffile', 'OpenCV', 'scikit-image', and 'ImageDraw' for image loading, format conversion, segmentation, contour detection, feature extraction, shape fitting, and mask generation across multiple microscopy formats.

**Scientific Computing:** 'NumPy' for array operations and vectorized calculations; 'SciPy' for distance transforms, filtering, morphological operations, and geometric algorithms; and 'math' for specialized trigonometric and geometric functions.

**Machine Learning and Neural Networks:** 'PyTorch' for deep learning computations, supporting SAM2 (Segment Anything Model 2) implementation for GPMV tracking; and 'Cellpose' for specialized GPMV segmentation with the 'cyto3' model.

**Data Analysis and Management:** 'pandas' for structured data organization, statistical operations, and CSV export; 'CSV' module for direct file operations; and 'json' for configuration storage and parameter serialization.

**Visualization:** 'matplotlib' for creating plots and interactive visualizations.

**User Interface:** 'tkinter' and 'ttk' for creating interactive interface components; and 'ImageTk' for integrating processed images within the GUI environment.

**System and File Operations:** 'os', 'pathlib', and 'shutil' for cross-platform file management and directory operations; 're' for regular expression pattern matching in filename parsing; and 'warnings' for diagnostic messages.

**Performance Optimization:** 'threading' and 'multiprocessing' for parallel execution of computationally intensive tasks; and 'tqdm' for progress monitoring and reporting during operations.

**Supplementary Video 1: GPMVs blebbing.** Vesiculation buffer was added to cells expressing Spike with soluble mCherry, and frames were captured at 30-second intervals using confocal microscopy. Magenta indicates the cytoplasm, blue indicates the nucleus, and white indicates the membrane. Scale bar: 10 μm.

**Supplementary Video 2: Time-lapse live confocal microscopy imaging of GPMV docking using the C-trap confocal microscope and micropipette aspiration.** A GPMV expressing ACE2-GFP (yellow) was trapped using a micropipette and brought close to a Spike-expressing GPMV (magenta) to observe docking dynamics. The time stamps (in seconds) indicate the progression from initial contact to docking, with frames captured at 1.016-second intervals. Scale bar 2 μm.

**Supplementary Video 3: Live confocal imaging of mobile GPMVs and AI-accelerated analysis.** The video displays a three-column presentation of a docking event: The left column shows live confocal microscopy of freely diffusing GPMVs captured at 1-second intervals, with an ACE2-GFP expressing GPMV (yellow) interacting with a Spike-expressing GPMV (magenta). The middle column demonstrates AI-based tracking of both vesicles over time, following their positions and movements. The right column shows automated interface detection that dynamically adapts to the changing orientations and positions of the two interacting GPMVs throughout the docking process. Scale bar 10 μm.

**Supplementary Video 4: Live confocal imaging of GPMVs triggering and fusion.** Two hours after mixing freely diffusing ACE2-GFP GPMVs with Spike-expressing GPMVs, when docking had reached a steady state, 0.0832 mM trypsin was added. Frames were captured at 700-millisecond intervals using confocal microscopy. Scale bar: 20 μm.

**Supplementary Video 5: Live confocal imaging of GPMVs fusion.** The video highlights three fusion events captured at 100 ms intervals. The left column shows composite images with time overlays, capturing the temporal dynamics of ACE2-GFP and Spike-expressing GPMVs during fusion. The middle column isolates the mCherry content (magenta), highlighting its diffusion from one GPMV to another during the fusion process.
